# Familial primary ovarian insufficiency associated with a *SYCE1* point mutation: Defective meiosis elucidated in humanized mice

**DOI:** 10.1101/2020.02.07.938639

**Authors:** Diego Hernández-López, Adriana Geisinger, María Fernanda Trovero, Federico F. Santiñaque, Mónica Brauer, Gustavo A. Folle, Ricardo Benavente, Rosana Rodríguez-Casuriaga

**Affiliations:** Department of Molecular Biology, Instituto de Investigaciones Biológicas Clemente Estable, Montevideo, Uruguay; Biochemistry-Molecular Biology, Facultad de Ciencias, Universidad de la República, Montevideo, Uruguay; Department of Genetics, Instituto de Investigaciones Biológicas Clemente Estable, Montevideo, Uruguay; Flow Cytometry and Cell Sorting Core, Instituto de Investigaciones Biológicas Clemente Estable, Montevideo, Uruguay; Department of Cell Biology, Instituto de Investigaciones Biológicas Clemente Estable, Montevideo, Uruguay; Department of Cell and Developmental Biology, Biocenter, University of Würzburg, Würzburg, Germany

**Keywords:** Idiopathic infertility, primary ovarian insufficiency, meiosis, synaptonemal complex, humanized mice

## Abstract

**Objective:** To investigate if nonsense mutation *SYCE1* c.613C˃T *-*found in women with familial primary ovarian insufficiency (POI)- is actually responsible for infertility, and to elucidate the involved molecular mechanisms.

**Design:** As most fundamental mammalian oogenesis events occur during the embryonic phase, thus hindering the study of POI’s etiology/pathogeny in infertile women, we have used CRISPR/Cas9 technology to generate a mouse model line with an equivalent genome alteration (humanized mice).

**Setting:** Academic research laboratories.

**Interventions:** We present the characterization of the biallelic mutant mice phenotype, compared to wild type and monoallelic littermates.

**Animals:** Studies were conducted employing the generated humanized mice. All studies were performed for both genders, except otherwise stated.

**Main outcome measures:** reproductive capability by fertility tests; gonadal histological analysis; evaluation of chromosome synapsis and synaptonemal complex (SC) assembly by immunolocalizations; protein studies by Western blotting; transcript quantification by RT-qPCR.

**Results:** The studied mutation proved to be the actual cause of the infertile phenotype, both in female and male mice homozygous for the change, confirming infertility of genetic origin with a recessive mode of inheritance. The mechanisms that lead to infertility are related to chromosome synapsis defects; no putative truncated SYCE1 protein was observed, and *Syce1* transcript was hardly detected in biallelic mutants.

**Conclusions:** We present for the first time the generation of humanized mice to study the actual consequences of a SC component mutation found in women with familial POI. By this approach we could confirm the suspected etiology, and shed light on the underlying molecular mechanism.

## Capsule

Humanized mice were generated to study the effects on fertility of a mutation in a synaptonemal complex-component-coding gene found in women with familial POI, enabling etiology confirmation and mechanism elucidation.

## Introduction

Primary ovarian insufficiency (POI) is a clinical syndrome characterized by loss of ovarian activity before the age of 40. It is a heterogeneous condition with a broad phenotypic spectrum, sharing the common feature of ovarian follicle dysfunction or follicle depletion. It can have serious noxious effects upon women’s psychological and physical health. POI incidence increases with age, affecting one in every 10,000 women at the age of 20, and 1 in 100 at the age of 40 [1]. Most cases (50–90%) have unknown causes and, therefore, are classified as idiopathic [2]. A significant contribution to idiopathic POI resides in the genetic background of the diagnosed females [3], with 10%–15% of them having an affected first-degree relative [4]. Among the already reported causes of POI, there are alterations in chromosome number and structure (e.g. Turner’s syndrome, 45,X), as well as genomic alterations in 46,XX non-syndromic patients [5–7]. During the last two decades, an increasing number of POI-associated genes have been identified both on the X chromosome [e.g. 8-11] and in autosomes [e.g. 8, 12-15], as well as in mitochondrial DNA [16], thus confirming the heterogeneous nature of the genetic causal component.

Given the requirement of meiotic divisions for normal gamete formation, it is expected that mutations in meiosis-related genes would account for at least part of the idiopathic infertility cases. Specifically, as due to their importance for recombination and proper chromosome segregation synaptonemal complexes (SCs) are essential structures for gametogenesis progression, alterations in SC-coding genes are obvious candidates to be at the groundwork of infertility [revised by 17], and particularly of POI. The SC is a meiosis-specific proteinaceous, ladder-like structure that physically binds together homologous chromosomes, and facilitates the resolution of recombination intermediates [18]. SCs are composed of two lateral elements (LEs), a central element (CE), and transverse filaments (TFs) linking both LEs with the CE. The CE together with the TFs constitute the SC central region (CR). So far, eight different SC protein components have been identified, including LE proteins SYCP3 [19, 20] and SYCP2 [21–23], TF constituent SYCP1 [24–26], and CE components SYCE1, SYCE2, SYCE3, TEX12, and SIX6OS [27–30].

The involvement of SC components in POI would be supported by loss-of-function studies for different SC genes employing KO mice, which have been reported to disrupt SC structure, and lead to infertility [22,25,29–35]. Some human mutations in SC-coding genes have been identified and linked to infertility [revision by 17]. Concerning LE components, various mutations for SYCP3 have been reported, and the first SYCP2 mutations have just been identified [36]. For some of the SYCP3 mutations, a dominant negative effect has been revealed in heterozygous individuals, in which the truncated SYCP3 interfered with polymerization of the normal protein [37, 38].

Regarding CE components of the SC, in the past years mutations potentially associated to clinical conditions have started to be reported. In particular, deletions in human 10q26.3 encompassing *SYCE1* gene were found in POI patients [39–41]. Besides, thus far three reports identifying mutations in *SYCE1* gene in infertile patients have been made [42–44]. In one of these reports, a homozygous point mutation was identified in a 13-member-family in which two sisters born to consanguineous parents suffered primary amenorrhea [42]. This mutation, *SYCE1* c.613C˃T, would lead to SYCE1 protein truncation. By sequencing studies, the authors determined that of the 11 descendants (five males and six females), only the two affected siblings were biallelic for *SYCE1* c.613C˃T mutation, suggesting a genetic cause with a recessive mode of inheritance [42]. Although the idea of a possible relation of SYCE1 mutation with pathogenicity would be supported by the phenotype of *Syce1* KO mice, which are infertile [35], an unequivocal evaluation linking SYCE1 mutations to the observed medical conditions is lacking so far.

As most fundamental mammalian oogenesis events (including SC formation and recombination) occur during the embryonic phase, eventual defects in this process are identified with many years of delay, leaving few possibilities to intervene, and even to study the condition’s etiology and pathogeny. A valid alternative to circumvent this difficulty is the employment of suitable animal models, which has the highest physiological relevance after human studies. However, thus far no transgenic humanized mice mimicking mutations found in humans for any SC component-coding gene have been reported.

In order to determine whether mutation *SYCE1* c.613C˃T is the actual cause of the observed infertile phenotype, and to study its pathogeny, we have generated a humanized mouse model line containing an equivalent point mutation *via* CRISPR/Cas9 mutagenesis system. Here, we present the phenotypic characterization of the humanized mutant mice, helping to shed light on the etiology and mechanisms of these infertility cases. We also discuss the potential usefulness of these humanized mouse models as substrates for future development of gene therapy approaches.

## Materials and methods

### Ethical approval

All animal procedures to generate the mutant line were performed at the SPF animal facility of the Transgenic and Experimental Animal Unit of Institut Pasteur de Montevideo. Experimental protocols were accordingly approved by the institutional Animal Ethics Committee (protocol number 007-18), in accordance with national law 18,611 and international animal care guidelines (Guide for the Care and Use of Laboratory Animals) [45]. All subsequent experimental animal procedures were performed at Instituto de Investigaciones Biológicas Clemente Estable (IIBCE, Montevideo, Uruguay), also in accordance with the national law of animal experimentation 18,611 (Uruguay), and following the recommendations of the Uruguayan National Commission of Animal Experimentation (CNEA, approved experimental protocol 009/11/2016).

### Design of molecules for mutagenesis and generation of humanized mice

CRISPR/Cas mutagenesis was employed aiming to obtain the desired humanized mouse cell line [42]. Design and selection of molecules for directed mutagenesis were carried out taking into account on-target ranking, off-target ranking, and distance of single-guide RNA (sgRNA) to target site of mutagenesis (http://www.broadinstitute.org/rnai/public/analysis-tools/sgrna-design). The selected sgRNA was acquired as *CRISPRevolution Synthetic sgRNA kit* (Synthego, USA). The single-stranded oligonucleotide donor (ssODN) employed as template for homology directed repair (HDR) was designed making use of online tools for silent mutation scanning (http://watcut.uwaterloo.ca/template.php) and restriction enzyme analysis (nc2.neb.com/NEBcutter2/), and ordered from IDT as 4 nmole Ultramer DNA Oligo (IDT, USA). Protospacer adjacent motif (PAM) was disrupted in the ssODN in order to avoid repeated nuclease action after the desired edition. Mice and zygote manipulation for genome editing was performed as previouly described [46, 47]. For details, see *Expanded discussion of the Materials and Methods*.

### Genotyping of transgenic mice

Offspring genotyping was performed *via* tail-tips. DNA was extracted by means of *GeneJET Genomic DNA Purification Kit* (Thermo Fisher Scientific, USA). The genomic region of interest (*i.e.* where the mutation was directed) was specifically amplified by standard PCR. The primers employed were: *Syce1*-613-FOR: 5’ TCAAGGAAGGTGAGGTCAGG 3’; *Syce1*-613-REV: 5’ ATGAAGAGACATACCGGCAG 3’. PCR products were run by electrophoresis, recovered by *GeneJET Gel Extraction and DNA Cleanup Micro Kit* (Thermo Fisher Scientific, USA), and sequenced.

### Fertility tests

Fertility was assessed both for females and males by mating 2-month-old mutant mice homozygous for the change with adult WT mice of opposite gender. Heterozygous mutants and WT mice were used as control groups. Assays were performed in triplicate for each gender in breeding pairs or trios (two females and one male). After at least 3 months without offspring, the analyzed individuals were considered infertile.

### Histology

Whole adult testes and ovaries were primary fixed in 2.5% glutaraldehyde, postfixed in 1% osmium tetroxide, dehydrated and resin-embedded (Durcupan, Fluka) according to conventional procedures [48]. Thereafter, 250 nm sections were cut using a *Power Tome XL* ultra-microtome (Boeckeler Instruments, USA), stained with toluidine blue, and examined by bright field microscopy. Photographs were taken by means of an *Olympus FV300* microscope equipped with a *DP70* camera, and *DPController v.1.1.1.65* software.

### Analysis by flow cytometry (FCM)

Testicular cell suspensions were prepared using a mechanical method previously described by our group [49, 50]. The resulting cell suspensions were stained with Vybrant DyeCycle Green (VDG, Invitrogen Life Technologies, USA) at a final concentration of 10 μM for 1 h at 35 °C in the dark with gentle agitation (80 rpm) as reported earlier [51]. FCM analyses were performed by means of a flow cytometer and cell sorter *MoFlo Astrios EQ* (Beckman Coulter, USA), using a 488 nm laser, a 100 μm nozzle (25 psi), and Summit software (Beckman Coulter, USA). For details concerning flow cytometer calibration and analyzed parameters, see *Expanded discussion of the Materials and Methods*.

### Antibodies

Primary anti-rabbit antibody against SYCE1 amino-terminal region (*i.e.* capable of detecting WT SYCE1 and its putative truncated form) was developed at GenScript (GenScript USA Inc.), using peptides ATRPQPLGMEPEGSC and CPEGARGQYGSTQKI from the amino-terminal region of the protein. This affinity-purified antibody was employed both for fluorescence microscopy (1:200) and for Western blots (0.3 µg/mL).

The other antibodies employed in this study were either commercial, or described in detail elsewhere [26]. For more information, see *Expanded discussion of the Materials and Methods*.

### Immunocytochemistry

Immunolocalization assays were performed on gonadal spread cells obtained through the dry-down technique [52] with minor modifications (see *Expanded discussion of the Materials and Methods*). For oocyte spreading, fetal ovaries (E18 embryos) were used, while for spermatocytes spreading we employed mechanically disaggregated adult testes. Slides were afterwards used for incubations with the indicated antibodies for immunofluorescence microscopy. All incubations with primary antibodies were performed overnight at 4°C in the presence of protease inhibitors (P2714, Sigma-Aldrich). Secondary antibody incubations were done at room temperature for 1 h protected from light.

### Microscopy and Imaging

All immunofluorescence microscopy acquisitions were performed employing a Zeiss LSM 800 confocal microscope (Carl Zeiss Microscopy, Germany) equipped with *Airyscan* processing module, a 63X/1.4 N.A. Plan Apochromat oil objective, Axiocam 506 color digital camera, and ZEN Blue 2.3 software (Carl Zeiss Microscopy, Germany). Airyscan image processing was done through the software’s automatic deconvolution step. All images were analyzed by means of FIJI ImageJ software [53].

### Statistical Analyses

Quantitative data of spread nuclei with synapsed chromosomes (zygotene and pachytene stage) from biallelic and monoallelic mutants with WT littermate controls were statistically compared using a chi-square test. Regarding RT-qPCR data, statistical significance and p-value were calculated in R bioconductor (http://cran.r-project.org/).

### Western blots

Testicular protein lysates corresponding to 7.5 x 10^5^ cells in Laemmli sample buffer were loaded per lane. SDS-polyacrylamide gel electrophoresis was carried out on 12% polyacrylamide gels. Protein gels were transferred to nitrocellulose membranes as instructed [54], and Western blots were performed as previously described [55]. Membranes were incubated for 2 h at room temperature in TBST with primary antibodies (anti-SYCE1-Nt and anti-β-tubulin), and for 1 h in blocking solution with anti-rabbit secondary antibody. Bound antibodies were detected by using the *Super Signal West Pico substrate* (Pierce). All assays were performed more than once, and using biological replicates.

### RT-qPCR assays

Total RNA from testicular cell suspensions was extracted with *PureLink RNA Mini Kit* (Ambion, Life Technologies, Carlsbad, CA), following manufacturer’s recommendations. RNA quantification was done by Nanodrop 1000 Spectrophotometer (Thermo Scientific). Retro-transcription and qPCR were performed using *Power SYBR Green Cells-to-Ct kit* (Ambion), starting from 50 ng of RNA, following kit instructions, in a CFX96 Touch Real-Time PCR Detection System 1 (BioRad, Hercules, CA). For qPCR step, 2 µL cDNA in 20 µL final volume reaction mix was used.

For RT-qPCR on embryonic ovaries, the same kit was directly employed after a lysis reaction with no previous RNA extraction (due to the scarcity of the tissue). The primers used are listed in Supplemental Table 1. We made 3 biological replicas, and chose *Ppp1cc* (*protein phosphatase 1, catalytic subunit, gamma isozyme*) as normalizing gene, as it has been previously shown to be a good normalizing gene for testicular RNA [56]. Amplification efficiency of all primers was >93%. The 2^-ΔΔCt^ method and WT mouse RNA as calibrator condition were used [57].

## Results

### Generation of model mouse line

Our first aim was to generate a model mouse line mimicking the *SYCE1* c.613C>T point mutation observed in humans. To achieve this, we chose the CRISPR/Cas9 technology, and proceeded as described in *Materials and Methods*. Comparison of SYCE1-coding regions from human and mouse genome evidenced high identity, thus facilitating the choice of the editing target (Fig 1A). *SYCE1* c.613C˃T is a nonsense mutation that would lead to a truncated human SYCE1 protein of 240 residues, while its WT counterpart has 351 amino acids. In mouse, an equivalent mutation would lead to a truncated protein of 242 amino acids, as compared to the WT 329-residue version.

**Figure 1.**
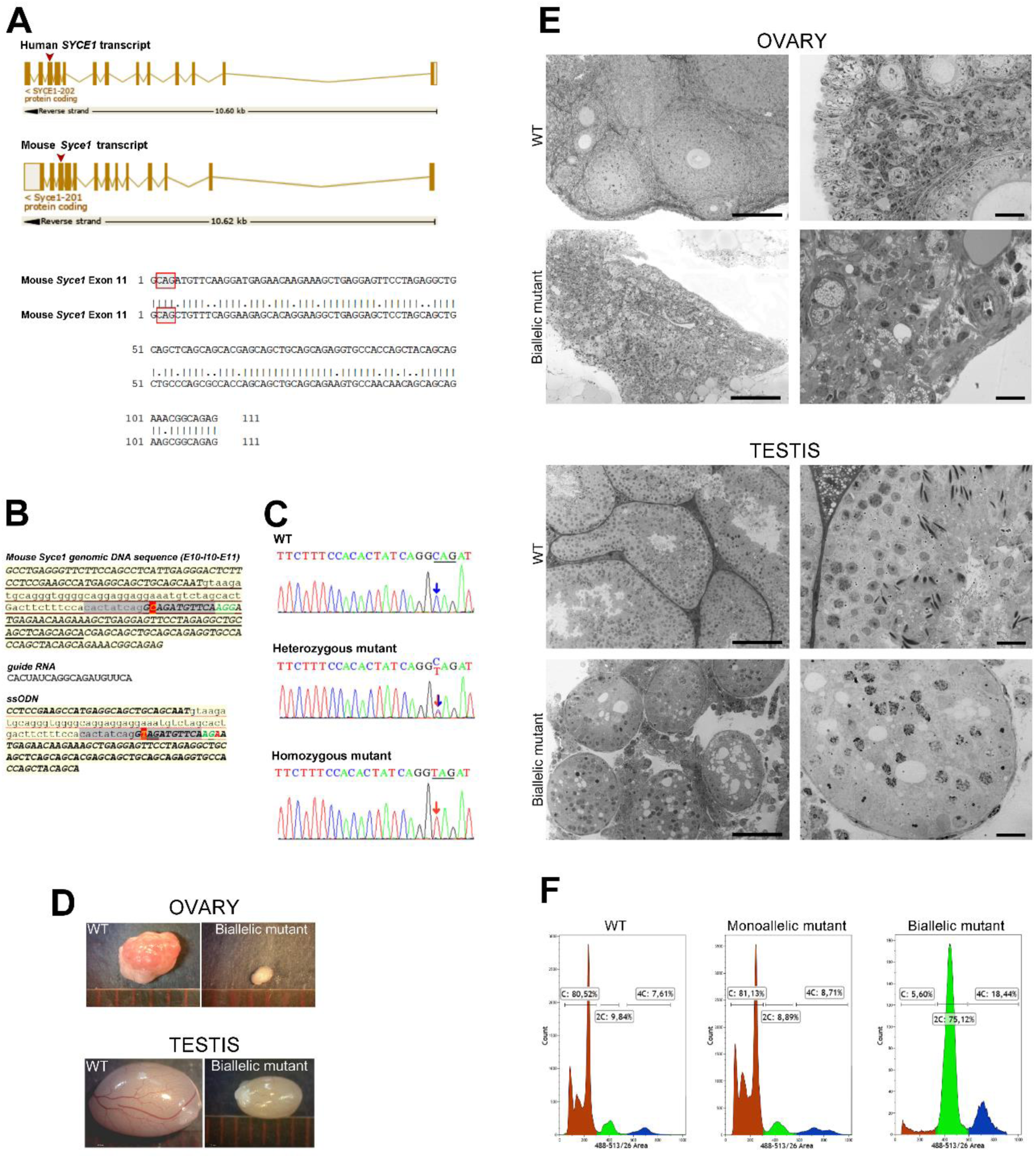
Mouse genome editing strategy and initial characterization of edited mice. (**A**) Graphical representation of *SYCE1/Syce1* transcripts from human and mouse. Boxes represent the 13 exons separated by intronic sequences (lines); arrowheads indicate exon #11 where the mutation was found in humans, and to which mutagenesis was directed in mouse genome. The alignment of mouse and human genomic sequence for exon #11, showing high sequence similarity, is also presented. Red boxes indicate the codon affected by mutation c.613C>T in humans that would generate a premature TAG stop codon. (**B**) Mouse *Syce1* genomic sequence corresponding to exon #10 + intron #10 + exon #11. The 20 nt sequence complementary to the sgRNA is shown in grey, protospacer adjacent motif (PAM) in green, and cytosine to be edited in red. Sequences of the sgRNA and single stranded oligonucleotide (ssODN) are indicated. Note in the latter the C>T substitution at the beginning of exon #11, as well as the disrupted PAM sequence. (**C**) Representative genotyping results obtained through standard sequencing of PCR products amplified from mouse tail-tips. (**D**) Comparative size of gonads in *Syce1* c.727C>T biallelic mutant and control mice. (**E**) Microscopic analysis of gonads in semi-thin sections of Epon-embedded ovaries and testes from adult WT and biallelic mutants. Panoramic view (*left*) and higher magnification images (*right*) are shown. Bars correspond to 100 and 20 µm (left and right images, respectively). (**F**) Flow cytometric analysis of testicular cell suspensions from adult WT and mutant mice. Representative FCM profiles obtained for testis from mutant mice and WT littermates are shown. Relative percentages of C, 2C and 4C cell populations are indicated in each case.

Design of molecules to be used in the directed mutagenesis (sgRNA and ssODN) was optimized to favor the HDR (homology directed repair) pathway (Fig 1B) [58]. Specimens resulting from microinjected zygotes (F0) were genotyped in search of the desired change, and then mated with WT individuals to obtain F1 generation. Afterwards, heterozygous specimens from F1 were intermated to generate F2 offspring. As expected, the latter included WT specimens as well as others heterozygous and homozygous for the desired point mutation, which in mouse corresponds to *Syce1* c.727C>T (Fig 1C).

### *Syce1* c.727C>T biallelic mutation causes infertility both in female and male mice

Fertility was assessed both for females and males by mating mutant mice with WT specimens of opposite gender. Data from three experimental groups was compared in these studies: WT, heterozygous and homozygous mice. No differences were observed between WT and heterozygous mice, which got as easily pregnant, and laid on average 7 pups with an equal ratio of male and female offspring. After 3 months, only individuals homozygous for the *Syce1* c.727C>T mutation failed to have offspring. This result was consistently reproduced in triplicates for each gender, and led us to conclude that the sole presence of this mutation in both alleles should be enough to produce the infertile phenotype observed in women [42]. Moreover, the same mutation in men should be able to cause infertility as well.

### *Syce1* c.727C>T biallelic mutation affects gonadal development

When gonadal size and aspect were assessed, no evident differences were found between adult WT and heterozygous mutant mice, neither for females nor for males (Supplemental Fig 1A). However, adult homozygous mutants showed striking differences in gonadal size and aspect as compared to their WT littermates, and this proved to be true both for female and male adult individuals (Fig 1D).

Concerning microscopic analysis of ovaries, growing oocytes and follicle development were evident in adult WT and heterozygous female animals (Fig 1E and Supplemental Fig 1B), while in their biallelic *Syce1* c.727C>T littermates no follicles or oocytes were observed (Fig 1E). Regarding testicular development, while both WT and heterozygous adult males showed normal seminiferous tubules with complete spermatogenesis (Fig 1E and Supplemental Fig 1C), the microscopic analysis of gonadal content from adult biallelic male mutant mice evidenced a severely affected spermatogenesis process with complete absence of post-meiotic stages (Fig 1E). The seminiferous tubules of these mutants were also depleted from mid and advanced prophase I stages (*i.e.* pachytene and diplotene), indicating an arrest in early meiotic prophase I stages. Moreover, the seminiferous tubules were much smaller than those of the WT and monoallelic mutants, and exhibited an unusual aspect (Fig 1E).

In order to have stronger quantitative comparative analyses, testicular cell suspensions from adult mice were analyzed by flow cytometry (FCM), mainly based on DNA content. Figure 1F shows representative FCM results. While no significant differences were found between WT and heterozygous mutants, this study confirmed for the biallelic mutant males complete absence of post-meiotic C population (Fig 1F). Regarding the 4C population, the FCM analyses hereby presented were obtained using the DNA-specific fluorochrome VDG that -as we had previously reported-allows the discrimination of two populations of spermatocytes: the early spermatocyte population (leptotene and zygotene stages, L/Z), and the mid/late spermatocyte one (pachytene and diplotene stages, P/D) [51]. Despite these populations are usually visualized in the histograms as a 4C bimodal peak (Fig 1F), this latter could not be observed in the FCM profiles from biallelic mutants (Fig 1F) that resembled those expected for 13-14 dpp WT juvenile mice [59].

### Evaluation of chromosome synapsis and SC assembly by confocal laser scanning microscopy

The dramatic effect of the point mutation on gonadal development, prompted us to study its consequences on SC structure and homologous chromosome synapsis. Immunocytochemical localizations were performed on spread cells from both embryonic ovaries and adult testes, and analyzed by confocal laser scanning microscopy.

A first set of studies was centered on SC lateral element component SYCP3, in order to evaluate chromosome synapsis. As the pachytene stage (completely synapsed homologs) is reached by day 13-14 post-partum in male mice and by day 17.5-18 post-coitum in female mice embryos, the ages of the specimens to be analyzed were chosen accordingly. Quantitative data of spread meiocyte nuclei with synapsed chromosomes from *Syce1* c.727C>T mutants and WT littermate controls were statistically compared using a chi-square test (*X*^2^). Once again, no evident differences were found between monoallelic mutants and WT littermates (*X*^2^ [1, *N* = 84] = 0.26, *p* = 0.61), with both presenting normally-looking spread chromosomes that had reached the pachytene stage (Fig 2A). However, biallelic mutants consistently showed impaired synapsis, presenting, at most, closely juxtaposed chromosomes that resembled earlier meiotic prophase stages (Fig 2A). These findings proved to be true for both genders (herein shown for females), and indicate that the sole biallelic presence of the point mutation severely affects homologous chromosome synapsis (*X*^2^ [1, *N* = 84] = 134.6, *p* < 0.00001), and would most probably account for the observed gametogenesis failure.

**Figure 2.**
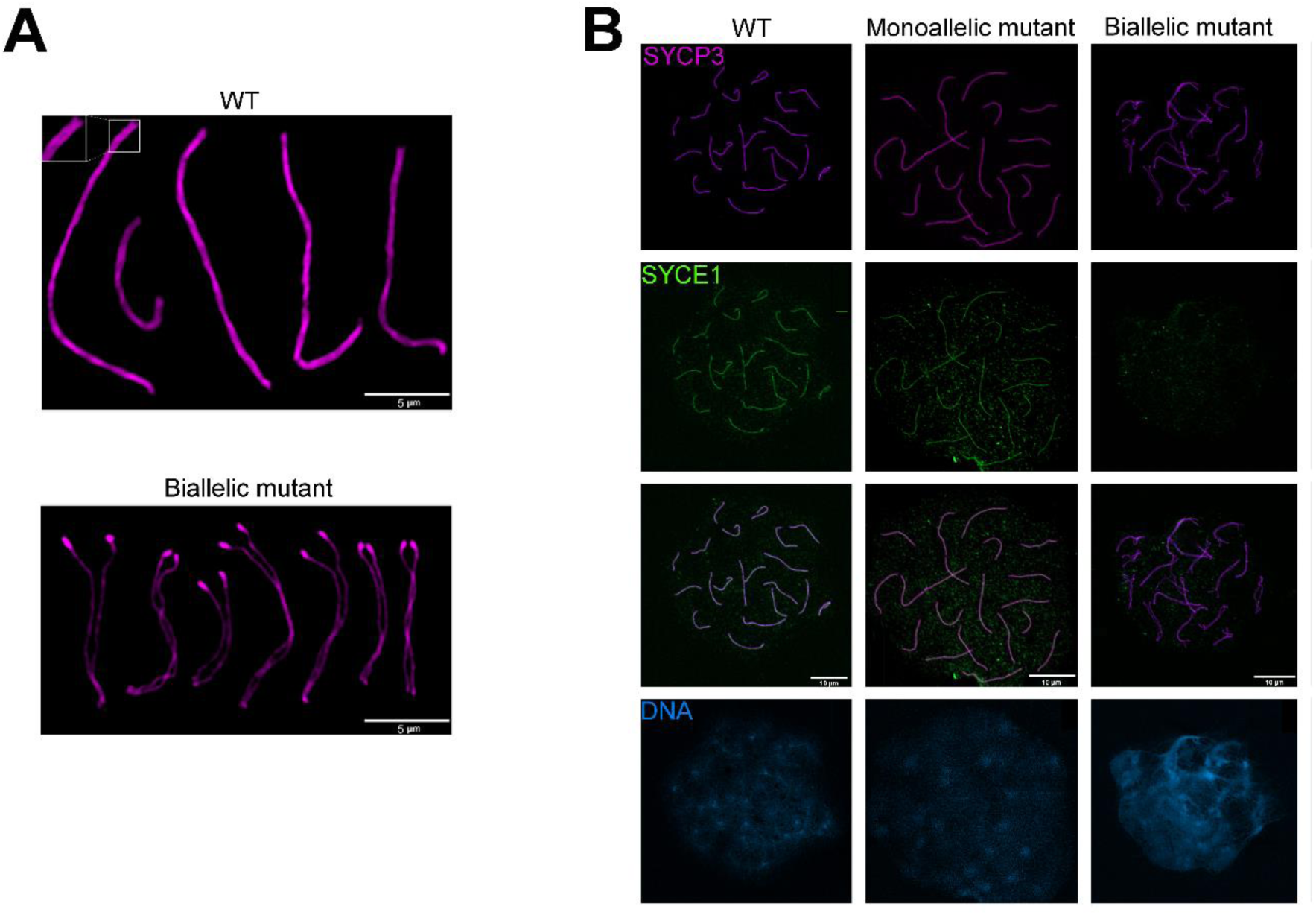
Evaluation of chromosome synapsis and SYCE1 loading to SC in WT and humanized mice. (**A**) Immunolabelling of LE component SYCP3 on spread meiotic chromosomes from WT (above) and biallelic mutant mice (below). Fluorescence acquisition was performed by means of an Airyscan module that enabled the resolution of LEs, even in completely assembled SCs (see inset above). Closely aligned but unsynapsed LEs are observed for biallelic mice (below). (**B**) Immunolocalization of SYCE1 protein in female 18 dpc WT and mutant mouse embryos. SYCE1 is shown in green, and SYCP3 in magenta. Merged channels and DNA staining with DAPI are also shown below.

As mentioned above, *Syce1* c.727C˃T is a nonsense mutation that would lead to a truncated protein of 242 amino acids, as compared to the WT 329-residue version. In order to evaluate the eventual presence of the putative truncated SYCE1 protein in the SC of mutants, we addressed protein immunolocalization employing an antibody specially developed against the SYCE1 amino terminal (N-t) region (see *Materials and Methods*). We clearly detected SYCE1 in spread meiocytes from WT and monoallelic mutant individuals, but not in those of biallelic mutants. This result was observed for either of both genders (herein shown for females; Fig 2B), and was consistently obtained for all biological and technical replicates.

Afterwards, we evaluated the presence of other known protein components of the SC central region (*i.e.* TFs and CE). Some of the components have been reported to be loaded earlier than SYCE1 onto the SC (*i.e.* TF SYCP1 and CE SYCE3), while others would be loaded later (*e.g.* CE TEX12) [22,28–30,60–64]. Representative results are shown in Figure 3. Again, no differences were found between WT and monoallelic mutants for any of the analyzed components either in female (Fig 3A-C) or male meiocytes (*e.g.* Fig 3D). Regarding biallelic mutants, protein components SYCP1 and SYCE3 were detected on spread meiocytes containing SCs in assembling process (Figs 3A,B), while TEX12 was not detected at all in the assembling structure (Figs 3C,D).

**Figure 3.**
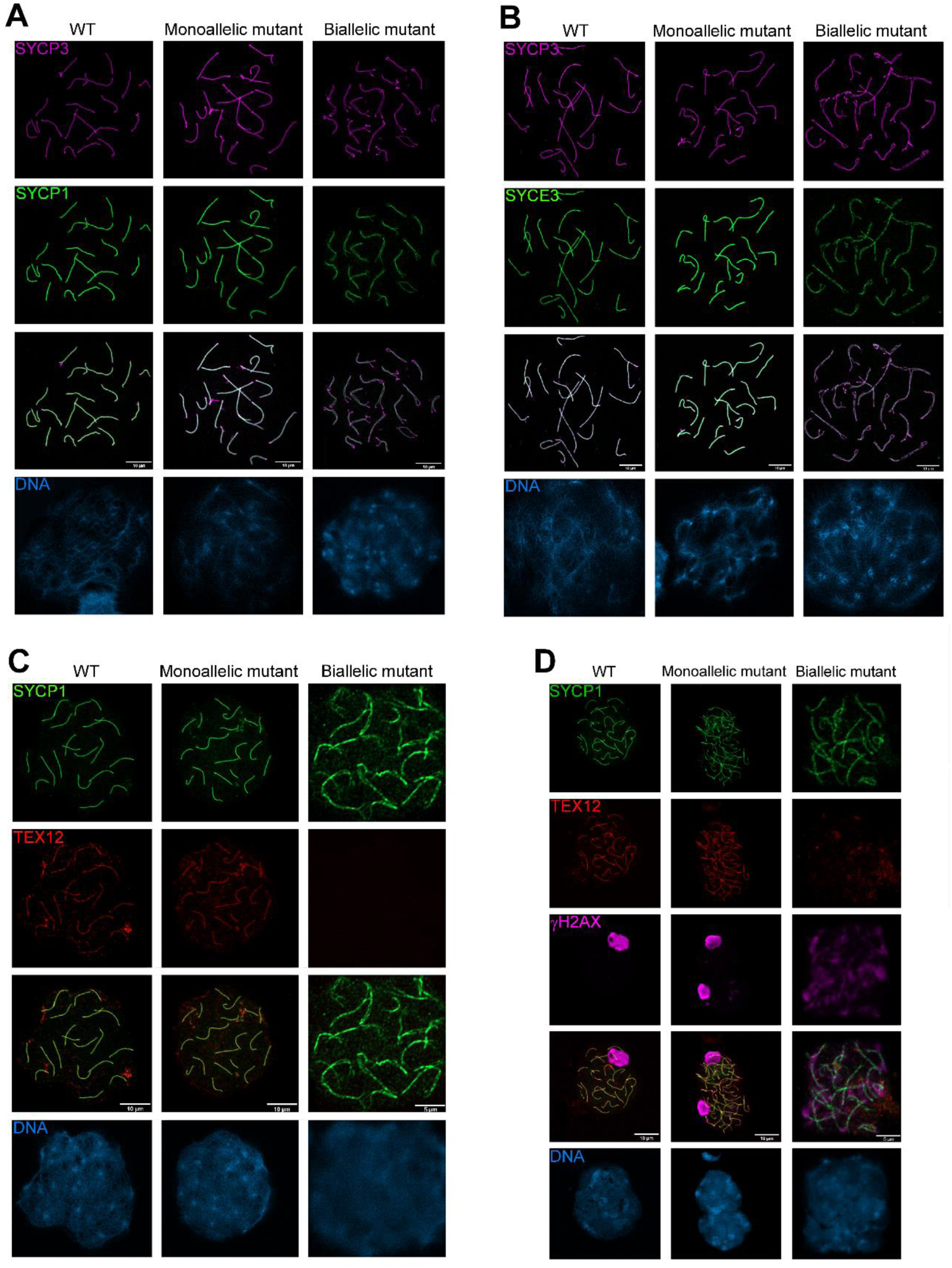
Immunolocalization of other CR SC components in WT and mutant mice. (**A**) Immunolabelling of TF SYCP1 in female 18 dpc mouse embryos. SYCP3 is shown in magenta, and SYCP1 in green. (**B**) Immunolocalization of CE SYCE3 in female 18 dpc mouse embryos. SYCE3 is shown in green, and SYCP3 in magenta. (**C**) Immunolabelling of CE TEX12 in female 18 dpc mouse embryos. SYCP1 is shown in green, and TEX12 in red. (**D**) Immunolocalization of TEX12, SYCP1 and γH2AX in male adult WT and mutant mice. SYCP1 is shown in green, TEX12 in red and γH2AX in magenta. Merged channels and DNA staining with DAPI are shown below in each case.

For spermatocytes, γH2AX was also immunolabeled along with SC protein components. This histone variant renders a very typical staining on male meiotic chromosomes: dispersed chromosome staining in early stages, then restricted to the XY body in pachytene stage. No difference in this regard was detected between WT and monoallelic male mutants (Fig 3D). However, as expected for a pre-pachytene meiotic arrest, no restricted staining for the sexual chromosome pair was found in biallelic male mutants, which presented a diffuse γH2AX staining pattern, characteristic of earlier meiotic stages (Fig 3D).

### The putative truncated SYCE1 protein is not detected in mutant mice testes

Although no SYCE1 protein was detected on assembling SCs of biallelic mutant mice, still, the putative SYCE1 truncated protein could be present in meiocytes, although not incorporated into the SC. In order to shed some light on the molecular mechanism leading to infertility, we assessed the presence of the putative truncated protein in mutant mice through Western blot analysis on testicular material. These protein studies cannot be performed in females due to material requirements unable to be fulfilled with embryonic ovaries (< 0.0001 g).

A band with an apparent molecular mass of 38 KDa was detected both for WT mice and monoallelic mutants, in accordance with the predicted molecular weight for WT 329-residue version of murine SYCE1 protein (Fig 4A). No truncated SYCE1 protein (theoretical expected size: 28 KDa) was detected for monoallelic mutants.

**Figure 4.**
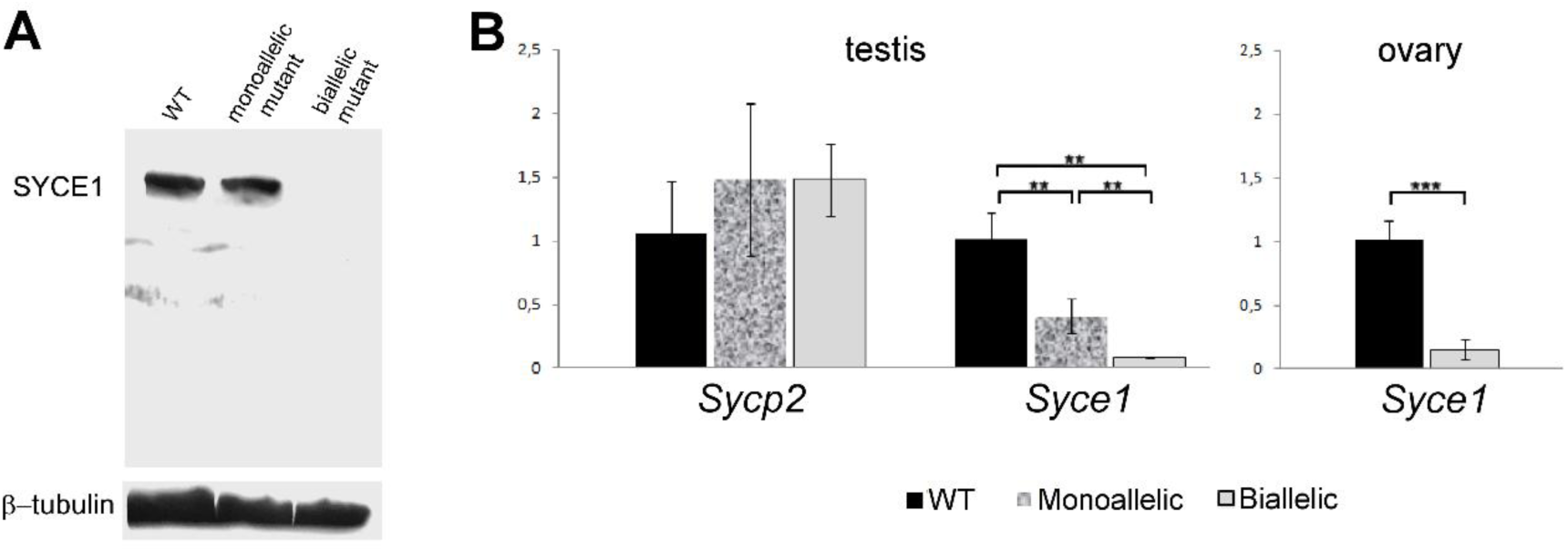
Analysis of SYCE1 protein by Western blot, and transcript quantitation by RT-qPCR. (**A**) Western blot analysis of SYCE1 protein and its putative truncated mutant variant. The blotted bands were immunodetected with a specific rabbit antibody against mouse SYCE1 Nt-region. β-tubulin was employed as loading control. (**B**) RT-qPCR results obtained for gonads from WT and mutant mice. Statistical levels of significance are indicated in each case. \*\**P value* < 0.005; \*\*\**P value* < 0.0001.

Concerning biallelic mutants, no protein reactive to SYCE1 antibody was detected at all (see Fig 4A). Protein gels were deliberately overloaded to minimize the effects of detection sensitivity limits, but in all assays no band of 28 KDa was observed.

### *Syce1* transcript is fairly detected in mutant humanized mice

The results from the Western blot assays prompted us to analyze transcript levels. As shown in Figure 4B, *Syce1* transcript quantitation rendered big differences between WT and biallelic mutants that exhibited minimum mRNA levels, both for embryonic ovaries (p< 0.0001) and for adult testes (p< 0.005).

Less pronounced but still significant differences were obtained between WT and monoallelic mutant males that showed intermediate transcript levels (p< 0.01; Fig 4B). On the contrary and as expected, quantitation of *Sycp2* transcript (coding for SC LE SYCP2) rendered no significant differences between the three conditions.

## Discussion

The present work deals with some primary ovarian insufficiency cases presumably related to mutations in SC coding genes, but still classified as idiopathic infertility. We have worked with nonsense mutation *SYCE1* c.613C˃T, addressing the question on its actual responsibility for the infertility observed in female patients homozygous for this mutation. We have also intended to shed some light on the underlying mechanisms involved. Aiming these, we have applied directed mutagenesis in mouse for genome editing, and successfully generated a humanized mouse line (*i.e.* the edited murine genome contains a mutation equivalent to the one found in humans). In this regard, the high percentage of identity between human and mouse SYCE1 coding genes allowed us to easily find a *SYCE1* c.613C˃T murine equivalent mutation (*Syce1* c.727C>T). Generation of this animal model has enabled us to study the etiology and pathogeny of these infertility cases.

Concerning etiology, it could be established that the biallelic presence of *Syce1* c.727C>T mutation is enough to cause infertility in both females and males. On the other hand, mice heterozygous for the mutation were as fertile as WT individuals. Thus, infertility in these cases has now proven genetic origin and recessive mode of inheritance. It is worth reminding that for humans, de Vries and collaborators (*i.e.* the authors that reported the mutation) found no clinical symptoms for heterozygous individuals of both genders examined in their study [42]. However, the phenotype for males homozygous for the mutated *SYCE1* gene could not be known at that time because all the males examined in that study were heterozygous for the mutation [42]. Thus, although the identification of this nonsense mutation was initially connected to cases of women infertility, we can now anticipate that the biallelic mutation would equally cause infertility in men.

The fertility results herein reported evidence an absence of sexual dimorphism for this CE-related mutation. This would be in accordance with previous observations for mice with loss-of-function of CR components-coding genes, which were equally infertile in both genders [25,29,30,33–35], as opposed to LE-component mutants that showed sexual dimorphism [22, 32].

Regarding gonadal development, no differences were found between monoallelic mutants and unaffected WT individuals of both genders. This is in accordance with the fertile phenotype observed in heterozygous humans, and also with our fertility tests results. However, analysis of gonads from adult biallelic mutant mice did show striking differences both for females and males: minute ovaries with absence of oocytes and follicles (indicative of POI), and testes with incomplete spermatogenic process, and 3-4 times smaller than in unaffected littermates. Female mutant mice bearing POI resemble the clinical description of the human sisters with biallelic *Syce1* c.613C˃T, ratifying the validity of the experimental model.

In males, the seminiferous epithelium of biallelic mutants showed not only absence of post-meiotic cells, but even of mid/late meiotic prophase stages, thus indicating an early meiotic arrest. We also analyzed testicular cell suspensions by FCM as this methodology represents a widely accepted means to analyze testicular cellular content, bearing very high quantitative analytical power and statistical weight. These analyses let us corroborate a severely affected spermatogenic process in biallelic mutants, with testes completely depleted from postmeiotic haploid cells with C DNA content (*i.e.* round and elongating spermatids, and spermatozoa). Moreover, concerning the 4C population (mainly composed of primary spermatocytes), FCM profiles obtained for biallelic mutants resemble those of ≈13 dpp mice that have not reached the pachytene stage yet [59]. This result is in agreement with the early meiotic arrest observed by microscopic analysis.

Analysis of spread chromosomes immunolabeled against SC protein components, coupled to the use of an *Airyscan* super-resolution module, enabled a detailed study of chromosome synapsis. This study revealed that the biallelic presence of *Syce1* c.727C>T severely affects homologous synapsis.

Regarding interactions between SYCE1 and other SC proteins, previous studies based on co-immunoprecipitation and yeast two-hybrid assays have identified interactions with SYCE3 and SIX6OS1 [29,30,62]. Besides, SYCE1 recruitment to the SC has been proposed to be mediated by SYCE3 [65]. Neither SYCE1 protein nor SYCE1-downstream-loading SC components (e.g. TEX12) could be detected on meiotic chromosome axes of biallelic mutant humanized mice. As the mutation under study is nonsense, two hypothesis concerning the pathogeny of these infertility cases arose: a) the putative SYCE1 truncated protein, which would lack 87 residues from the carboxyl terminus (Ct), would not be recruited and loaded to the SC as its interaction with SYCE3 would be impaired, thus affecting normal SC assembly after SYCE3 loading step; b) impaired loading could be due to the absence of the putative truncated SYCE1 protein.

In order to shed some light on this matter, SYCE1 protein was assessed by Western blot assays. Even in overloaded protein gels, we have not been able to detect the putative truncated protein in the biallelic mutant mice. Similarly, in monoallelic mutant mice only the WT form of the protein could be detected. Thus, the impaired synapsis phenotype observed in biallelic mutant mice would result from the absence (or at least presence of undetectable levels) of truncated SYCE1, supporting our second hypothesis. Of note, the absence of an interfering shorter version of SYCE1 could explain the unaffected phenotype of monoallelic mutants. This would be quite different from the case of some *SYCP3* mutations, where a dominant negative effect has been reported in heterozygous patients [37, 38]. The lack of a possibly interfering truncated protein also has important implications concerning the development of eventual therapeutic procedures in biallelic patients, as it would guarantee the occurrence of no relevant interference with an eventually introduced exogenous normal protein, thus facilitating the intervention.

As the mutation studied herein is a nonsense one, a possible explanation for the lack of detectable mutant protein could rely on nonsense-mediated mRNA decay (NMD). This regulatory pathway functions to degrade aberrant transcripts containing premature termination codons (PTCs). Since mutations that generate PTCs cause approximately one-third of all known human genetic diseases [66], NMD has been proposed to have a potentially important role in human disease. *SYCE1* c.613C>T mutation could be one of these cases. In order to have a primary evaluation of this possibility, we performed *Syce1* transcript quantitation in humanized mice compared to WT littermates. The results obtained were consistent and pointed to transcript degradation of aberrant transcripts, with hardly detectable levels of transcript in biallelic mutant gonads of both genders (Fig 4B). In addition, monoallelic mutant males showed intermediate *Syce1* levels, in accordance with NMD pathway involvement. Thus, the biallelic presence of *Syce1* c.727C>T mutation would lead - through a different mechanism - to a similar phenotype to that reported for Sy*ce1 KO* mice, in which complete absence of SYCE1 protein causes infertility [35].

The findings herein reported represent a proof of principle, since there are no previous reports on the employment of CRISPR/Cas technology to direct a specific change to a SC component, and generate a humanized mouse model line for its exhaustive study. Besides, the generated mouse model line can be further employed in other studies, including those aiming to develop eventual therapeutic procedures.

## Acknowledgements

The authors thank the staff of Transgenic and Experimental Animal Unit core facility from Institut Pasteur de Montevideo, Uruguay, for technical support in the generation of the GM model.

## Expanded discussion of the Materials and Methods

### Mice manipulation for genome editing

Mice were housed in individually ventilated cages (Tecniplast, Milan, Italy), in a controlled environment at 20 ± 1°C with a relative humidity of 40-60%, in a 14/10 h light-dark cycle. Autoclaved food (Labdiet 5K67, PMI Nutrition, IN, USA) and water were administered ad libitum.

Cytoplasmic microinjection was performed in C57BL/6J zygotes using a mix of 20 ng/µL sgRNA, 30 ng/µL Cas9 mRNA, and 20 ng/µL ssDNA oligo. The same day, surviving zygotes were transferred to B6D2F1 0.5 dpc pseudopregnant females (25 embryos/female in average), following surgery procedures established in the animal facility [46]. Previously, recipient females were anesthetized with a mixture of ketamine (100 mg/kg, Pharmaservice, Ripoll Vet, Montevideo, Uruguay) and xylazine (10 mg/kg, Seton 2%; Calier, Montevideo, Uruguay). Tolfenamic acid was administered subcutaneously (1 mg/kg, Tolfedine, Vetoquinol, Madrid, Spain) to provide analgesic and anti-inflammatory effects [47]. Pregnancy diagnosis was determined by visual inspection by an experienced animal caretaker two weeks after embryo transfer, and litter size was recorded on day 21 after birth.

### Flow cytometry analysis

Flow cytometer calibration and quality control were carried out using *3.0 µm Ultra Rainbow Fluorescent Particles* (Spherotech, USA). Fluorescence emitted from VDG was detected with a 513/26 bandpass filter. The following parameters were analyzed: forward scatter (FSC-Height with P1 Mask), side scatter (SSC-Height), 513/26-Area (VDG fluorescence intensity), and 513/26-Width. Doublets were excluded using dot plots of 513/26 pulse-area vs 513/26 pulse-width. FCM data was analyzed with Kaluza software (Beckman Coulter, USA).

### Cell spreading

Fetal ovaries were dissected, incubated in hypotonic buffer (30 mM Tris-HCl pH 8.2, 17 mM sodium citrate, 5mM EDTA, 50 mM sucrose, 5mM DTT) for 30 minutes, mechanically disaggregated on clean slides containing 100 mM sucrose, fixed in 1%-paraformaldehyde/0.15%-TritonX100, and allowed to dry slowly (overnight in closed humidity chamber, then open). Once completely dry, slides were wrapped in aluminum foil, and stored at -80°C until use. For spermatocytes spreading, the same procedure was applied on mechanically disaggregated adult mice testes.

### Antibodies

Guinea pig anti-SYCP3 (1:200), guinea pig anti-TEX12 (1:200), rabbit anti-SYCP1(1:200) and rabbit anti-SYCE3 (1:200) primary antibodies were used as affinity purified immunoglobulins and described in detail elsewhere [26]. Mouse anti-γH2AX was purchased at Millipore (1:500, 05-636; Millipore, Germany). Primary anti-β-tubulin antibody employed in Western blots as loading control- was acquired from Abcam (ab6046, 1:8,000, Abcam Antibodies), and revealed using an anti-rabbit secondary antibody coupled to horseradish peroxidase (1:30,000, Pierce).

Suitable secondary antibodies coupled to AlexaFluor dyes were acquired from Invitrogen Life Technologies, USA: AlexaFluor488 goat anti-rabbit (A11034, 1:1,000), AlexaFluor633 goat anti-guinea pig (A21105, 1:1,000), AlexaFluor546 goat anti-guinea pig (A11074, 1:1,000).

## Supplemental material

**Supplemental Figure 1.**
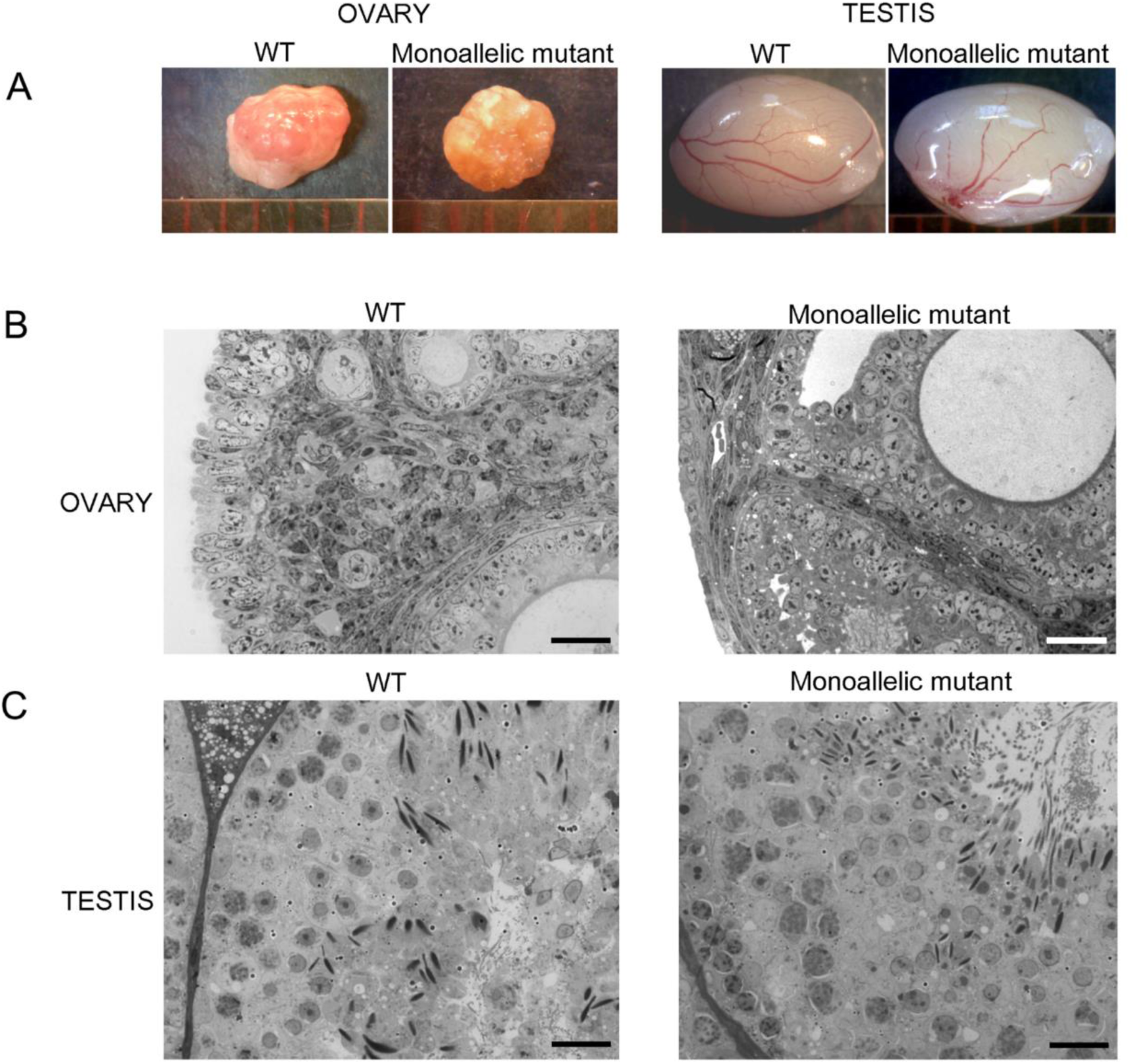
Macroscopic and microscopic analysis of gonads in WT and *Syce1* c.727C>T monoallelic mutant mice. (**A**) Comparative size of gonads. (**B**) Semi-thin sections of Epon-embedded ovaries from adult WT and monoallelic mutants. Note the presence of normal developing follicles both in WT and heterozygous mutant females. (**C**) Cross sections of seminiferous tubules from adult WT and monoallelic mutant mice. Normal spermatogenesis is evident in both cases. Bars correspond to 20 µm.

**Supplemental Table 1.**
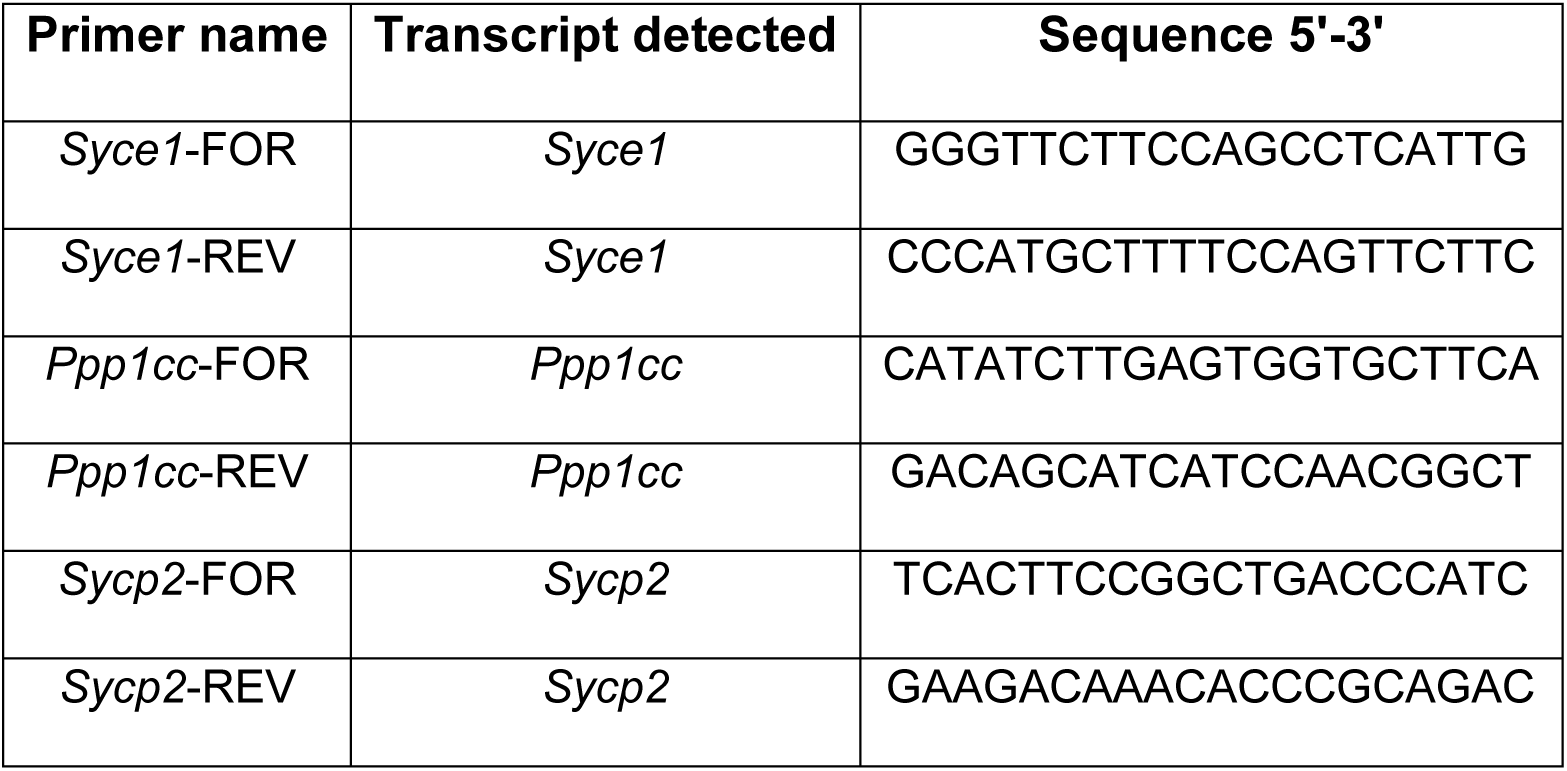
Primers used for qPCR experiments.

